# An integrated transcriptomics and proteomics study of Head and Neck Squamous Cell Carcinoma – methodological and analytical considerations

**DOI:** 10.1101/024059

**Authors:** Anupama Rajan Bhat, Manoj Kumar Gupta, Priya Krithivasan, Kunal Dhas, Jayalakshmi Nair, Ram Bhupal Reddy, HV Sudheendra, Sandip Chavan, Harsha Vardhan, Sujatha Darsi, Lavanya Balakrishnan, Shanmukh Katragadda, Vikram Kekatpure, Amritha Suresh, Pramila Tata, Binay Panda, Moni A Kuriakose, Ravi Sirdeshmukh

## Abstract

High throughput molecular profiling and integrated data analysis with tumor tissues require overcoming challenges like tumor heterogeneity and tissue paucity. This study is an attempt to understand and optimize various steps during tissue processing and in establishing pipelines essential for integrated analysis. Towards this effort, we subjected laryngo-pharyngeal primary tumors and the corresponding adjacent normal tissues (n=2) to two RNA and protein isolation methods, one wherein RNA and protein were isolated from the same tissue sequentially (Method 1) and second, wherein the extraction was carried out using two independent methods (Method 2). RNA and protein from both methods were subjected to RNA-seq and iTRAQ based LC-MS/MS analysis. Transcript and peptide identification and quantification was followed by both individual-ome and integrated data analysis. As a result of this analysis, we identified a higher number of total, as well as differentially expressed (DE) transcripts (1329 vs 1134) and proteins (799 vs 408) with fold change ≥ 2.0, in Method 1. Among these, 173 and 86 entities were identified by both transcriptome and proteome analysis in Method 1 and 2, respectively, with higher concordance in the regulation trends observed in the former. The significant cancer related pathways enriched with the individual DE transcript or protein data were similar in both the methods. However, the entities mapping to them were different, allowing enhanced view of the pathways identified after integration of the data and subsequent mapping. The concordant DE transcripts and proteins also revealed key molecules of the pathways with important roles in cancer development. This study thus demonstrates that sequential extraction of the RNA and proteins from the same tissue allows for better profiling of differentially expressed entities and a more accurate integrated data analysis.

**Author Contributions:** ARB, MKG, PK and SK contributed final data analysis. KD and JN were involved in the RNASeq experiments while MKG, SHV LB and SC were involved in the iTRAQ MS/MS analysis. RBR and HV contributed towards the standardization of sample collection and processing, and were also involved in obtaining clinical information of the patients along with SD. VK and MAK were involved in study design, providing clinical insights into the analysis and in critical assessment of the manuscript. ARB, MKG and PK were involved in manuscript preparation. AS, PT, BP, MAK and RS were involved in the establishing the study design, overall monitoring of the experimental results and manuscript preparation. PT, MAK, BP and RS are the lead investigators of the project.

**Significance of the study:** The study highlights the need to optimize tissue processing and analytical pipelines to enable accurate integrated analysis of high throughput omics data; a sequential extraction of RNA and protein entities and subsequent integrated analysis was identified to provide a better representation of the molecular profile in terms concordant entities and pathways.

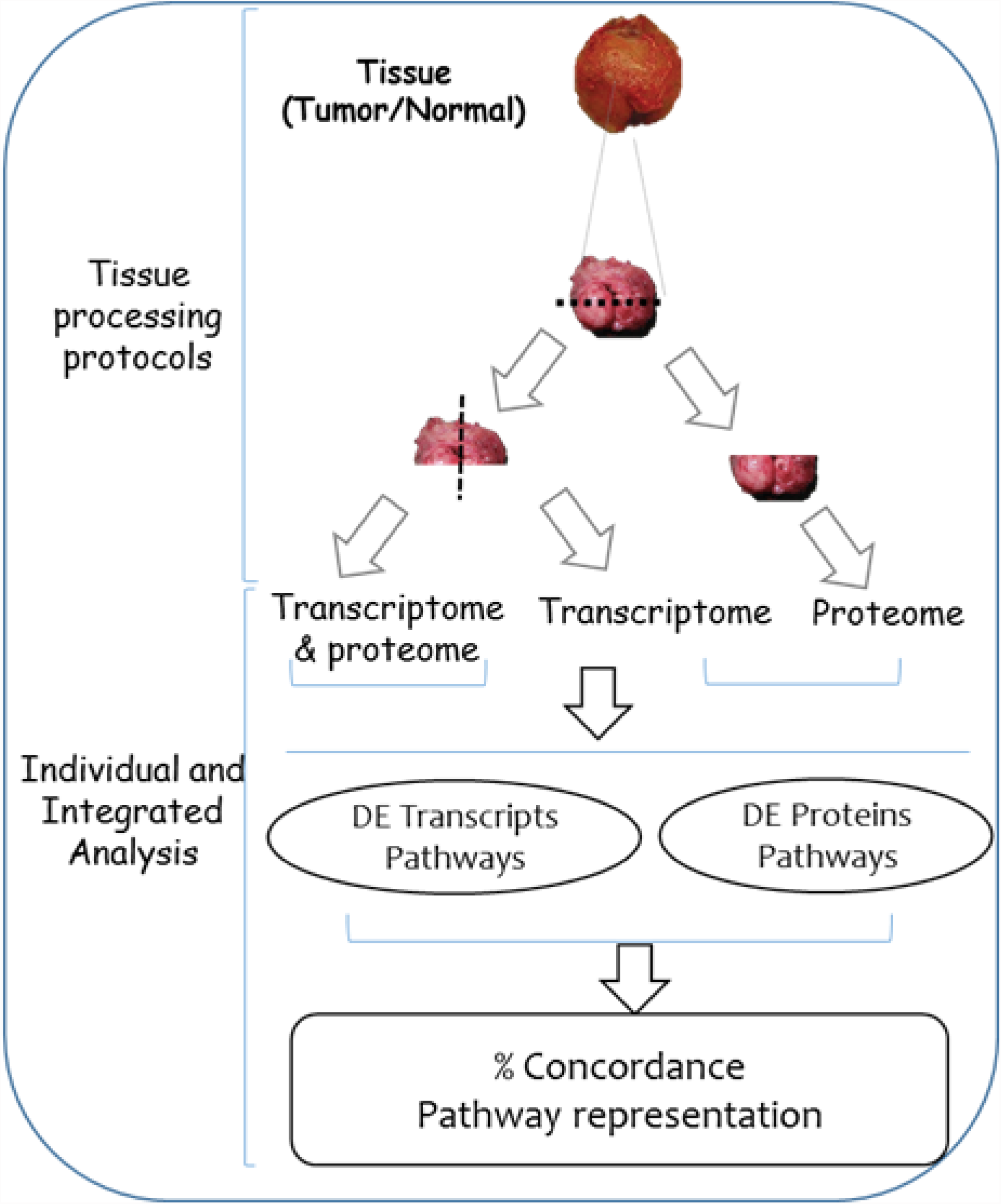

## Introduction

High-throughput omics approaches and data integration are being used to develop biomarkers for early detection or precision therapy. Intra-tumoral heterogeneity has been documented for several cancers and is a major challenge in biomarker identification and also an important consideration in integrated analysis ^1-4^. Intra-tumor heterogeneity evolves over time as the tumor progresses leading to spatial differences in genomic and molecular architecture in tumors. This phenomenon is reported in many solid tumors including head and neck squamous cell carcinoma (HNSCC) ^5-7^. Therefore, strategies to minimize the variables arising from intra-tumor heterogeneity are fundamental for integrated analysis of differentially expressed transcripts and proteins. In the current practice, the tissue samples used for transcriptomic and proteomic analyses are generally subjected to protocols wherein RNA and protein are extracted from different parts of the tumor. These could possibly represent spatially and molecularly different tumor regions, and hence introduce larger variations or discordance at transcript and protein level ^8, 9^. Therefore, optimal extraction protocols that enable uniform extraction of multiple analytes from the same biopsy samples may be useful. This is particularly important in many sub-sites of head and neck cancer, such as cancers of the larynx and pharynx with varying cyto-architechture of the biopsy tissue and also its limited availability for such analysis.

In this study, we generated transcriptomic and proteomic profiles from the surgical biopsy specimens of laryngo-pharyngeal cancers in order to evaluate the choice of the extraction protocol and to understand the important analytical considerations encountered during the process of integration of the two datasets for biological interpretation.

### Materials and Methods

A brief outline of the methods used is given here with the supporting details provided in the **Supplementary file 1**

The study was approved by the Institutional Review Board and the Institutional Ethics Committee. Informed consent was obtained from the participating subjects. Surgical tumor specimen and adjacent normal were collected from two patients with cancers of the larynx and pharynx. These samples from each patient were divided into two portions, one of which was stored in RNAlater for extraction of RNA alone, and both RNA and protein simultaneously and the other portion flash frozen immediately for protein extraction by sonication (**Figure 1A**).

**Figure 1.**
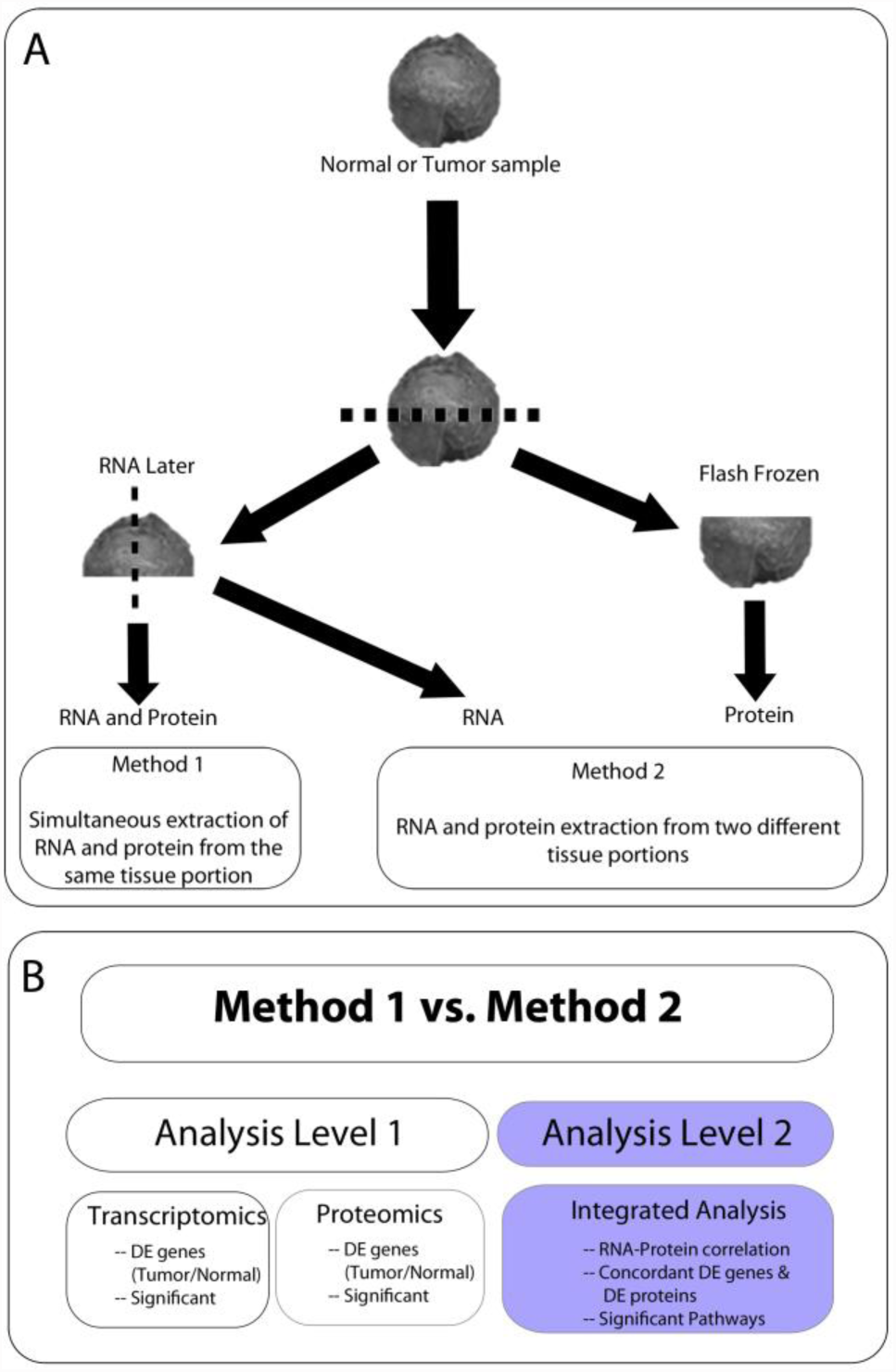
**Schematic of tissue processing and analytical pipelines (A)** Schematic diagram of isolation protocol. The tissue samples of both the normal and tumor of the same patient’s RNA and protein were isolated with two different protocols. Samples stored in RNA later and snap frozen samples were utilized to carry out the RNA and protein isolation. Two different methods were involved in the isolation of RNA and Protein; simultaneous isolation of RNA and protein (Method I) from the same chunk of tissue samples and independent isolation from different tissue portions of the same sample by different methods (Method II). The samples isolated by both the methods were used for downstream processing. **(B)** The isolated samples from the above mentioned methods were used for two different analysis; Level 1 involved independent transcriptomics and proteomics analysis and Level 2 refers to the integrated analyses of both the omics data. The differentially expressed (DE) gene entities were identified from Level 1 while the RNA-Protein correlation, significant pathways and concordant DE genes and DE proteins were assessed at Level 2.

Two different methods were used to isolate RNA and protein. In first method (Method 1) RNA and protein were extracted sequentially from the same tissue portion using NucleoSpin RNA/Protein kit (Macherery Nagel) as per manufacturer’s instructions detailed in supplementary file 1. In the second method (Method 2), RNA and protein were Isolated separately from different portions of the surgical specimen. The tissue stored in RNAlater was used for isolating RNA using Purelink RNA isolation kit (Ambion) and the flash frozen tissue was used for isolating protein in 0.5% SDS by sonication method.

The isolated RNA samples were quantified using Qubit RNA assay kit and their integrity assessed using RNA Nano 6000 chip on bioanalyzer instrument (Agilent, 2100). The RNA samples were then subjected to RNA sequencing protocol using the Illumina system HiSeq 2000 system after multiplexed cDNA libraries were prepared using TruSeq RNA Library preparation kit. RNA sequencing data was analyzed in Strand NGS software (version 2.1 Build 212125 © Strand Genomics, Inc., San Francisco, CA, USA). The DESeq2 algorithm proposed by Love *et* al^10^ was used to identify significantly differentially expressed transcripts (FDR 5%) between the tumor and normal samples in each method, considering the two patients as replicates. Relevant reads, genes and regions of interest from Strand NGS were subsequently imported into GeneSpring 13.0 (Agilent Technologies) to allow identification of enriched pathways from the Kyoto Encyclopedia of Genes and Genomes (KEGG) database^11^. Differentially expressed (DE) transcripts between the tumor and normal exhibiting a fold change ≥ 2.0 were used for all comparative analysis.

The quantification of protein was done by using Pierce™ BCA Protein Assay Kit. 100μg equivalent proteins from the four samples from one patient (Method 1-Normal and Tumor; Method 2 - Normal and Tumor) was used for the trypsin digestion followed by multiplex iTRAQ labeling according to manufacturers’ procedure (**Supplementary file 1**). The labeled peptides were pooled, vacuum-dried and reconstituted and fractionated by strong cation exchange (SCX) chromatography using an Agilent 1200 series LC system. Each fraction was then subjected to LC-MS/MS analysis using LTQ-Orbitrap Velos mass spectrometer (Thermo Fischer Scientific, Bremen, Germany) coupled with Proxion Easy nLC system (Thermo Scientific, Germany). The entire procedure was repeated for the other patient.

Proteome Discoverer Beta Version 1.4 (Thermo Fisher Scientific Inc., Bremen, Germany) was used for the database searches. MS/MS data was searched using SEQUEST search algorithm against the RefSeq protein database (version 65) and proteins were identified with 1% FDR at peptide level. Protein identification with 2 unique peptide(s) was considered for further analysis. Relative quantitation of proteins was estimated based on the relative intensities of reporter ions released during MS/MS fragmentation of their peptides. Differentially expressed (DE) proteins between the tumor and normal exhibiting a fold change ≥ 2 were used for all comparative analysis. All identified proteins were imported into GeneSpring 13.0 for pathway analysis. The differentially expressed transcripts and proteins were subjected to further bioinformatics analyses as described under results.

### Results

We have chosen two methods of RNA and protein extraction and applied the protocols to two sample pairs of patients diagnosed with laryngopharyngeal cancer. From each patient, the analytes were extracted from tumor and adjacent normal tissues for differential expression analysis. In one protocol, referred to as Method 1, RNA and protein extraction were carried out from the same aliquot of the homogenized tissue (tumor or surgical normal), while in Method 2, RNA and protein extractions were performed independently from two different portions of the tissues. We present here the analysis of the molecular profiles obtained from two patients using these two methods and present the scope of data integration.

The results from the two methods were compared in two stages; in the first stage, they were compared at the transcriptomic and proteomic level separately, for the coverage and then for DE entities. In the second stage, the RNA and protein data from each of the two methods were integrated and compared for their concordance in the differential expression. Concordant entities were considered for comparison of the two methods and also for mapping to pathways. In both stages, a pathway analysis was carried out to identify enriched KEGG pathways in GeneSpring 13.0 to compare the two methods at the functional level. The Multi-Omics Analyses workflow in GeneSpring was used to generate an enhanced pathway view with a simultaneous overlay of both transcript and protein data. The entire analysis pipeline is shown in **Figure 1B**.

### Analysis Stage 1

#### Transcriptome analysis

All the RNA samples were assessed for their quality to ascertain their compatibility for the analysis. The coverage statistics for each sample are shown in the supplementary document (**Table S1**). A total of approximately 20,000 transcripts were identified in each method. 1329 and 1134 transcripts were DE in the tumor samples, in Method 1 and Method 2, respectively. All the identified DE transcripts were significant (p-value ≤ 0.05) in both methods with a fold change ≥ 2.0. 851 DE transcripts were common to both methods, with all of them showing concordance in their regulation trends, i.e., they were identified as upregulated or down regulated in both methods **(Figure 2A & B)**.

**Figure 2.**
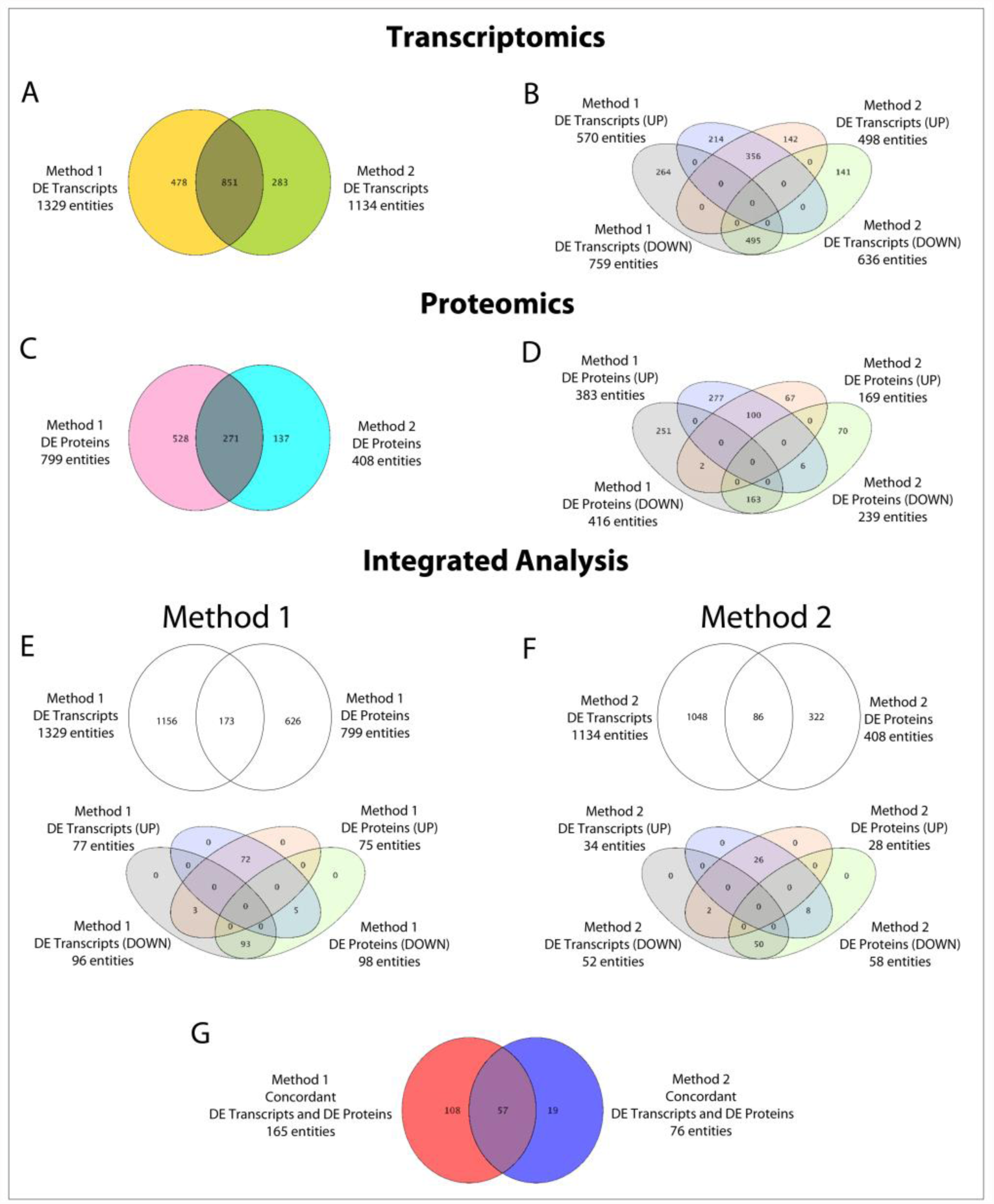
**Identification of differentially expressed (DE) RNA and protein between the Methods** A comparison between the transcriptomic analysis revealed a total of 851 gene entities between the two methods (a), the details of concordant entities (in terms of regulatory trends –UP or DOWN regulated) has also been provided (b). Comparison between proteomics datasets revealed higher differentials (N=528) in Method I than Method II (N=137) (c). The concordant gene entities with similar regulation trends is represented in the figure (d). The Integrated analysis of DE transcripts to proteins revealed a higher concordance in the Method I (N=173) (e) as compared to Method II (N=86) (f). 57 concordant entities were common between the two methods (g).

Of all the pathways enriched by DE transcripts from each of the two methods (p ≤ 0.05), we selected 20 top pathways commonly linked to cancer and found 15 to be common between the two methods. Cell cycle process, ECM-receptor interaction, PI3K-Akt Signaling Pathway, Glutathione Metabolism, and Focal Adhesion figured as the most significant pathways enriched in both the methods (**Figure 3A**). The number of genes mapping to each of these pathways were also comparable across the two methods.

**Figure 3.**
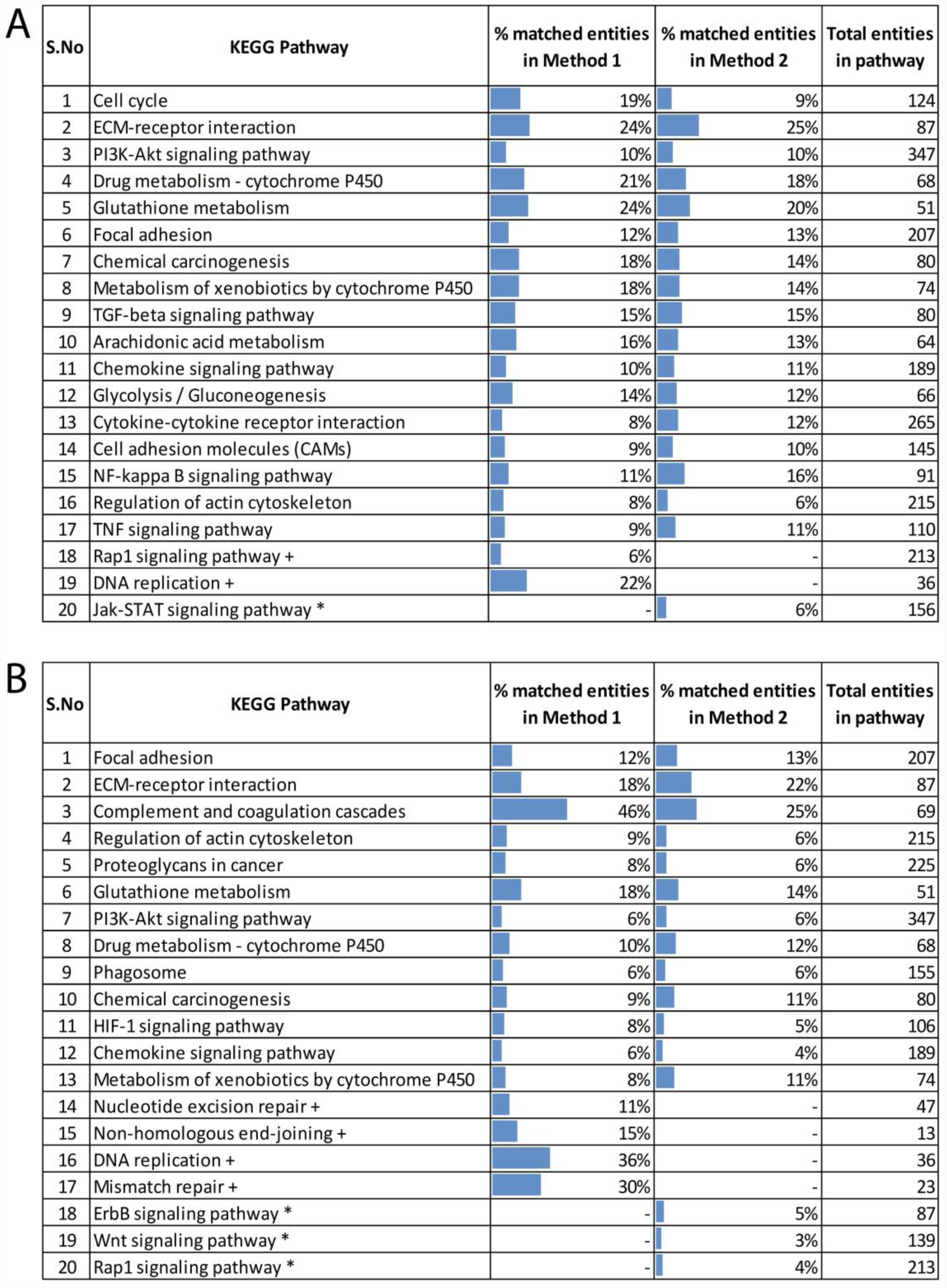
**Representative list of pathways relevant to cancer** Significantly enriched pathways identified by DE transcripts (A) and DE proteins (B) are represented. The pathways that are not significant in Method 1 (*) or in Method 2 (+) are indicated in both the figures.

#### Proteome Analysis

Proteins extracted from tumor and normal tissues using both the methods were subjected to trypsin digestion followed by iTRAQ labeling and LC-MS/MS analysis for protein identification and quantification. For proteins, the peptide level results from the two patients were integrated to generate a non-redundant dataset of protein identifications. About 3700 proteins, each with at least 2 unique peptides were identified in both the methods. Of these, 799 and 408 proteins were differentially expressed with fold change ≥ 2 in Method 1 and 2, respectively. 271 DE proteins were common between the two methods with 97% of them showing concordance in their regulation trends, i.e., they were upregulated or downregulated in both methods (**Figure 2C & D**). Thus Method 1 uniquely yielded many more proteins compared to Method 2.

We mapped the DE proteins to KEGG pathway and compared 20 top pathways commonly related to cancer, and found 50% of them to be common between the two methods. The number of proteins mapping to these pathways was also comparable. Further, most of these pathways were same as the ones identified at the transcript level that included ECM-receptor interaction, Focal Adhesion, PI3K-Akt Signaling Pathway and Glutathione Metabolism (**Figure 3B**).

Thus, Analysis level 1 showed that Method 1 resulted in a higher number of differentially expressed genes and proteins as compared to Method 2. However, at each omics level, a majority of DE entities and top pathway representations were comparable between the two methods. The specific entities mapping to them were however, not the same at the transcript and protein levels, thus allowing merging of the two type of entities to enhance the pathway view (see below).

### Analysis level 2

#### Integrated Analysis

The integration of transcript and protein data suffers from one challenge, in that the number of proteins accessed with the present analysis platforms is much smaller than the number of transcripts. Therefore, any integration is limited by the total number of proteins accessed and number of DE proteins observed. Genes differentially expressed at both RNA and protein levels were identified for integrated analysis by using Entrez Gene ID as the common identifier. In Method 1, 173 genes were differentially expressed proteins overlapped with transcripts (95% (n=165) being concordant) while in Method 2, 86 proteins had corresponding transcript matches (88% concordant, n=76) (**Table 1, Supplementary files 2 & 3).** The Spearman rank correlation between RNA and protein was higher in Method 1 (ρ = 0.77) as compared to Method 2 (ρ = 0.55).

Thus, in both the methods the number of genes showing concordance at both transcript and protein level was relatively small compared to the total number of DE transcripts and DE proteins. Method 1 scored higher even in this respect. Although significant pathway enrichment analysis with such small concordant transcript and protein datasets would be limiting, we could identify many molecules that are functionally important in HNSCC including IGFBP, ERK, COX2, STAT, PFN2, EPCAM, SERPINH1 and MCM2 and genes involved in angiogenesis, prolactin signaling and DNA repair.

The data integration may, however, be applied to enhance the strength of the entities mapped to pathways with individual -ome analysis. As discussed above, the pathways enriched by the DE transcripts and proteins observed in the two methods were comparable. However, the specific transcript or protein entities mapping to them were not the same, thus allowing development of an integrated view of the pathway by overlaying all DE entities identified at the two levels onto the pathways.

Phosphatidylinositol-3-kinase/protein kinase B (PI3K-Akt) signaling pathway, one of the most significant pathways in HNSCC, can be used as an example to look into the coverage of the different entities. In this study, pathway analysis revealed that the PI3K-Akt signaling pathway was enriched independently with both transcriptome and proteome datasets (**Figure 3)**. Higher number of differentials at each -omics level in Method 1 is manifested as greater number of matches in pathways (**Figure 4A & B**).

**Figure 4.**
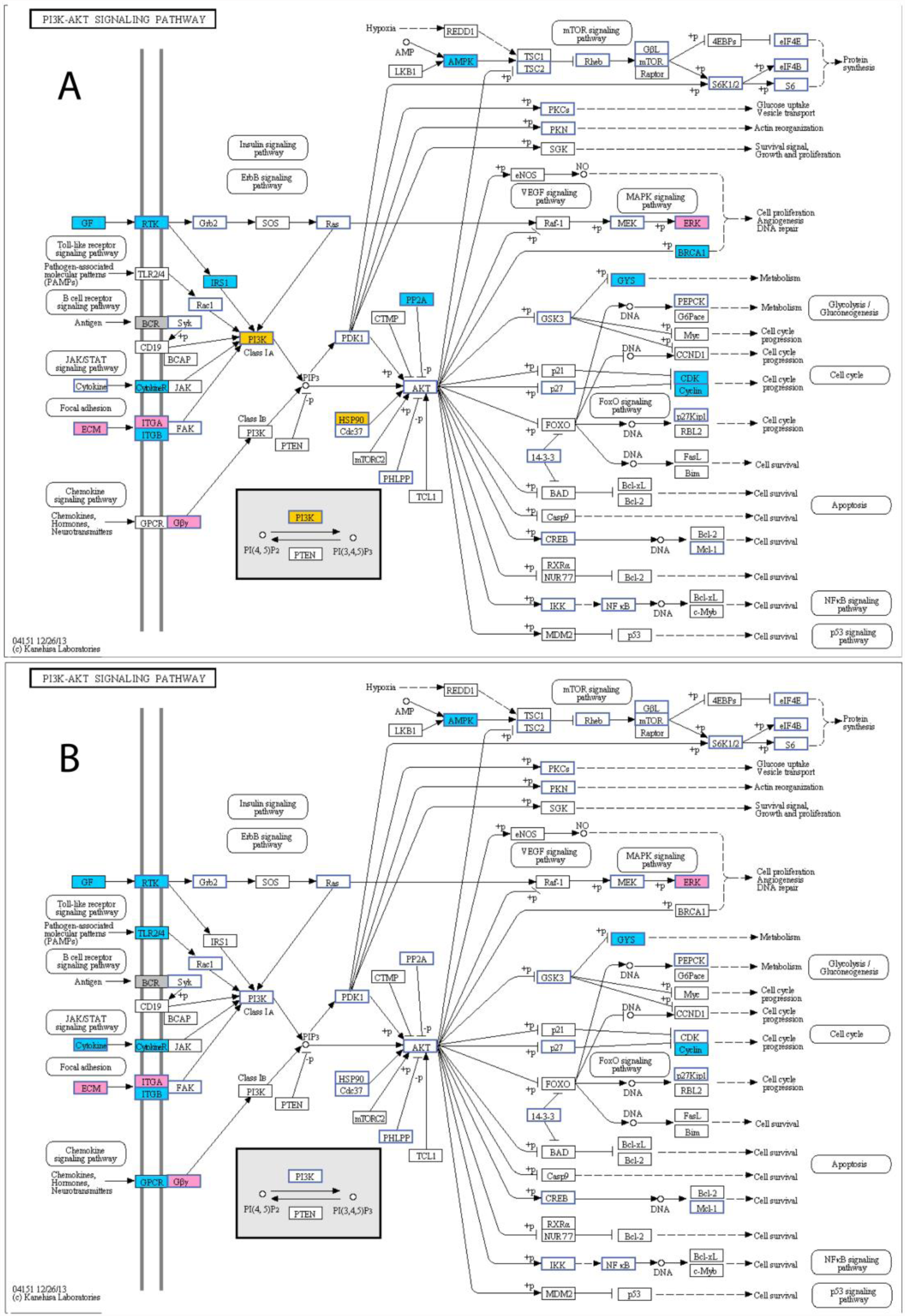
**Pathway representation of the differentially expressed transcript and protein entities** PI3K-Akt signaling pathway represented by matches from all DE transcripts and DE proteins in (a) Method 1, and (b) Method 2.

### Discussion

Integrated analysis of multi-omics data and pathway based approaches are being explored for identification of tumor-specific gene signatures, prediction of response to therapy, tumor recurrence and metastasis to improve diagnosis and design better therapies in cancer^12-16^. Spatial heterogeneity, an inherent property of several solid tumors, may cause multiple sections of the tumor biopsies to differ from each other, both with respect to expression as well as mutational profiles can have confounding effects on interpretation of multi-omics integrated data. Sample paucity is another factor which challenges optimal integrated analyses pipelines. To address these issues for multi-omic analysis of laryngo-pharyngeal cancer, we compared two different RNA and protein extraction methods; simultaneous extraction of both molecules from same tissue section (Method 1) and separate extractions from two different portions of the biopsy (Method 2). The methods were compared within the perspective of integrated analysis of multi-omic datasets.

Analysis of the individual transcriptomic and proteomic data using two methods, showed higher number of the DE entities identified in Method 1, suggesting that this method might provide a better global profiling. Integrated analysis is currently limited by the lower coverage of proteins in the proteomics experiments. Nevertheless, it has a significant value in the context of strengthening the identification of functionally relevant transcripts/proteins, through overlaying the two datasets. In this study, the 20% of the DE proteins were covered by the DE transcripts, which is consistent with the overlap generally observed^17-19^. This number reduced further when concordant DE transcripts and proteins were examined from the total matches. The technical or biological reasons underlying the discordance cannot be generalized but may be assessed at individual gene/protein; some possibilities could be differences in RNA and protein stability, post-transcriptional regulation, or translational efficiency^17, 20-22^. Further, it is also possible that the analytical stringency applied for differential expression at RNA and protein levels and concordance may not necessarily reflect the actual biological context. It should also be noted that the concordant matches would be higher if the fold change thresholds are made more flexible to accommodate even values that are close to but not exactly 2-fold change used as threshold. Nevertheless, a higher number of concordant DE transcripts and proteins was revealed in Method 1 when compared to Method 2.

Overlaying the data obtained by enrichment of pathways and processes by the individual - ome data offered higher coverage of the pathway. The PI3K pathway, one of the most important signal transduction pathways in cancer, overexpressed in 50-80% of HNSCCs^23, 24^, is one of the top pathways enriched with both individual transcript and protein data, and was used to illustrate the above point.. The PI3K-AKT pathway regulates a number of cellular processes including apoptosis, proliferation, cell cycle progression, cytoskeletal stability and motility, and mutations/amplifications of members of this pathway are commonly studied in cancers^24^. Identification of this pathway at the individual ‘omics’ level and its enhancement after merging the two data greatly increases the confidence of the analysis.

Integrated analysis of the individual omics data also identified many molecules relevant to the carcinogenic process in HNSCC: Mimecan preproprotein (OGN) identified by both the methods has been identified as prognosticator in laryngeal cancer while others such as KRT17, PFN2 have been reported to be dysregulated in HNSCC^25, 26^. ERK, COX2 and STAT1 also have been reported to play an important role in HNSCC progression and development of chemoresistance, and have also been proposed as potential targets for effective therapy ^27-31^. Integrated analysis of transcripts and proteins from Method 1 further identified unique molecules such as EPCAM, SERPINH1 and MCM 2. Epithelial cell adhesion molecule (EpCAM) is known to be involved in migration, cell proliferation and differentiation in HNSCC^32^ while Minichromosome maintenance complex (MCM2) and SERPINH1 were known to be predictive of malignant progression and poor prognosis^33^. Thus, an integrated analysis adds further value to single omics analysis by its ability to extract complementary information. Further, it may also be useful in identifying ‘high confidence’ leads to prioritize follow up and validation.

Overall, qualitative and quantitative assessments of the two methods show that sequential extraction of RNA and protein from the same tissue (Method 1), offers some advantages.

1. It requires smaller amount of the clinical tissue sample, is technically compatible for simultaneous isolation and downstream processing of two analytes including the chemistry used for quantitative proteomic analysis, and thus maybe the method of choice for integrated analysis
2. Although the pathway representation with the two datasets was comparable for both the methods, Method 1 resulted in higher numbers of DE transcripts and proteins mapping to any one pathway, in comparison to Method 2.
3. Method 1 also yielded higher numbers of concordant DE transcripts and DE proteins on integration of transcriptome and proteome data.

Our study thus provides a basic framework of analysis of tumor tissues using transcriptome and proteomic data and their integration. Although both transcriptomics and proteomics technologies have advanced significantly during the past decade, the depth of the proteome achievable in routine analysis is still much smaller that of the transcriptome. Current advancement in proteomics do offer prospects of deeper access to the proteome including protein variants. With these developments, the spectrum of integration of the omics data at these two levels would certainly expand further. There are also other possibilities not shown here, such as assigning isoforms not known at the protein levels by integrating RNA sequence data through a proteo-genomic approach, and mapping to peptides which remain unassigned due their absence in the libraries.

## Acknowledgements

The authors acknowledge the financial support from Department of Biotechnology Government of India for the project [BT/01/CEIB/11/IV/05 Dated: 22/08/13].

## Tables

**Table 1.** Overlap between DE transcripts and DE proteins in each method

## Supplementary Files and Tables

**Supplementary file 1:** Detailed Materials and Methods

**Supplementary Table 2 A:** List of overlapping differentially expressed transcript and protein entities of Method 1.

**Supplementary Table 2 B:** List of concordant differentially expressed transcript and protein entities of Method 1.

**Supplementary Table 3 A:** List of overlapping differentially expressed transcript and protein entities of Method 2.

**Supplementary Table 2 B:** List of concordant differentially expressed transcript and protein entities of Method 2.

**Supplementary table 1:** Coverage statistics (total number of aligned reads) for each sample

## Detailed Materials and Methods

### Clinical sample collection

The study was approved by the Institutional Review Board and the Institutional Ethics Committee. Informed consent was obtained from the participating subjects. Surgical tumor specimen and adjacent normal were collected from two patients with laryngo-pharyngeal cancers. The treatment naïve samples (larynx and hypopharynx) were collected from patients who were both from male (42-60 years), with risk habits and diagnosed with moderately differentiated squamous cell carcinoma. These samples from each patient were divided into two portions, one of which was stored in RNAlater and another flash frozen immediately after procurement. In Method 1, a portion of the tissue stored in RNAlater was used to extract both RNA and protein simultaneously while in Method 2, a portion of the tissue in RNAlater was used for RNA extraction while the flash frozen tissue was used to isolate protein (**Figure 1**).

### Materials

The molecular biology chemicals used in the study were procured from different sources; RNAlater (Ambion); NucleoSpin RNA/Protein Kit (Macherey-Nagel, USA); ß-mercaptoethanol and SDS (Sigma Aldrich); Molecular Grade ethanol (MERCK); Purelink RNA isolation kit (Ambion, USA); Qubit RNA assay kit (Invitrogen, USA); TruSeq RNA Library preparation kit (TruSeq RNA Sample Prep v2, Illumina, USA); HS DNA bioanalyzer kit (Agilent, 2100); Qubit HS kit (Invitrogen, USA); TruSeq SBS kit v3 chemistry; Pierce™ BCA Protein Assay Kit; Trypsin (Promega, Trypsin Gold – Mass Spectrometry grade V5280, Rev 3/13); iTRAQ labels (Applied Biosystems, iTRAQ® Reagents Multiplex Kit, Version 5.2 Revision Date 30.01.2013).

### Method 1: Extraction of RNA and protein simultaneously from the same tissue portion

Approximately 50mg of both tumor and normal tissue samples stored in RNAlater were used for the isolation of RNA and protein from the NucleoSpin RNA/Protein Kit (Macherey-Nagel, USA). 350μL Buffer RP1 and 3.5 μL ß-mercaptoethanol (ß-ME) was added and homogenized using a tissue homogenizer. This was further used for the combined isolation of RNA and protein as per the manufacturer’s instruction. RNA bound to the column is eluted using RNase free water. The flow through containing the protein was precipitated, the pellet washed with 50% ethanol, dried for 15min in RT and later dissolved in 0.5% SDS. The RNA and protein isolated was stored -80°C till further analysis.

### Method 2: Extraction of RNA and protein separately from different tissue portions

The tissues (∼10mg) (normal and tumor) stored in RNAlater were used for isolation of RNA using Purelink RNA isolation kit (Ambion, USA). The tissues were lysed and homogenized with lysis buffer containing 1% β-mercaptoethanol, vortexed and 1 volume of 70% ethanol was added to the lysate and vortexed for thorough mixing. The lysed samples were transferred to the spin column and centrifuged at 12000rpm at RT for 1min, the flow through was discarded and the column was washed as per recommendations using the wash buffer containing ethanol. Elution was done in 30ul of RNAse free water. The isolated RNA samples were quantified using Qubit RNA assay kit (Invitrogen, USA) and their integrity assessed using RNA Nano 6000 chip on bioanalyzer instrument (Agilent, 2100). Both the RNA samples were then subjected to RNA sequencing protocol using the Illumina system.

### Protein Extraction by sonication method

Approximately ∼50mg of tissue was washed thoroughly in 1% PBS and 100μL of 0.5% SDS was added and sonicated for 10 seconds for 7-8 times each. This was then centrifuged at 14000rpm at 4°C for 30min and the supernatant was collected. The protein estimation was done by BCA method. The sample quality and the normalization was checked by running a normal SDS-PAGE in a medium gel.

### Transcriptomics and Proteomics

### RNA Library preparation and RNA sequencing

Multiplexed cDNA libraries were prepared using TruSeq RNA Library preparation kit (TruSeq RNA Sample Prep v2, Illumina, USA) following manufacturer’s instructions for the RNA extracted for tumor biopsies from two patients along with surgical normal margin, using two methods. 500ng of total RNA was used for library preparation. The final libraries were checked for quality and quantity using HS DNA bioanalyzer kit (Agilent, USA) and Qubit HS kit (Invitrogen, USA) respectively. Libraries were sequenced using 2x100 bp paired-end TruSeq SBS kit v3 chemistry following manufacturer’s instructions. Sequencing was carried out using the RNA-seq sequencing protocol in the Illumina HiSeq 2000 system.

### Quantitative LC-MS/MS analysis

Quantitative proteomics for differentially expressed proteins in the tumor samples as compared to the surgical normal margins was carried out by iTRAQ labelling of tryptic peptides followed by mass spectrometry analysis.

#### a. Trypsin digestion, iTRAQ labeling and fractionation of peptides

Hundred-microgram equivalent proteins from the four samples from one patient (Method 1-Normal and Tumor; Method 2 - Normal and Tumor) was used for the trypsin digestion followed by iTRAQ labeling according to manufacturers’ procedure. In brief, to each of the four samples, 20μL of dissolution buffer and 1μL of reducing agent was added and incubated at 60°C for 60min. Further, the cysteine blocking reagent (2μL) was added to these samples and incubated for 10min at RT. The protein samples were then trypsinized using 4μg sequencing grade trypsin (1:25) for 16h at 37°C. The tryptic peptides were then labeled with 4-plex iTRAQ reagents which dissolved in 70 μl of ethanol at room temperature for 1 h. The reactions were quenched with equal volume of water. Sample labeling was as follows: peptide samples from Method 1 (Normal and Tumor tissue samples) with 116 and 117 tags, respectively and peptide samples from Method 2 (Normal and Tumor) with 114 and 115 tags, respectively. The labeled peptides were then pooled, vacuum-dried and reconstituted in 10 mM KH_2_PO_4_, 20% acetonitrile (pH 2.8) (solvent A) and fractionated by strong cation exchange (SCX) chromatography (PolySULFOETHYL A column, 100 × 2.1 mm, 5 μm particles with 300 Å pores; PolyLC, Columbia, MD) using an Agilent 1200 series LC system. A gradient of 50 min from 5 to 40% Solvent B (350 mM KCl in solvent A) with flow rate of 300 μl/min was applied. A total of 15 fractions were obtained and desalted using stage-tips packed with C18 material. The samples were then vacuum dried and taken further for LC-MS analysis. The entire procedure was repeated for the other patient.

#### b. LC-MS/MS

Each peptide fraction was reconstituted in 0.1 % TFA and subjected to LC-MS/MS analysis using LTQ-Orbitrap Velos mass spectrometer (Thermo Fischer Scientific, Bremen, Germany) coupled with Proxion Easy nLC system (Thermo Scientific, Germany). In house Magic C18 AQ reversed phase material (Michrom Bioresources, 5 μm, 100 Å) was used to make chromatographic capillary columns. Peptide samples were enriched using Trap column (75 μm × 2 cm) at a flow rate of 3μL/min followed by separated by analytical column (75 μm × 10 cm), at a flow rate of 350 nL/min. A linear gradient of 7-30% ACN was used for 60 min to elute the peptides. Data was acquired in a data dependent mode at a mass resolution of 60,000 at 400 m/z. High-energy collision dissociation (HCD) was used for fragmentation at 41% collision energy with a dynamic exclusion window of 45 seconds. Top twenty peptides were selected for fragmentation from each duty cycle for MS/MS and detected at a mass resolution of 15,000 at m/z 400. The automatic gain control for full FT MS was 1 million ions and for FT MS/MS was set to 0.1 million ions with a maximum time of accumulation of 250 milliseconds. The lock mass option was used to increase accuracy of mass measurements.

### Data Analysis

### Transcriptome data: Transcript identification and quantitation

RNA sequencing data was analyzed in Strand NGS software (version 2.1 Build 212125 © Strand Genomics, Inc., San Francisco, CA, USA). Reads from each sample were aligned against the Human hg19 build from UCSC using the Strand NGS proprietary aligner. The alignment was carried out with a minimum match length of 25, minimum percent identity of 90 and a maximum of 5 gaps allowed. Pre and post alignment QC was done to assess data quality based on base, read and tile characteristics. Reads that failed standard Illumina QC metrics, average base quality less than 20 or alignment score less than 98 were filtered out. Aligned reads were further quantified against the RefSeq (version 62) gene model and normalized across all samples in a method specific manner. The DESeq2 algorithm proposed by Love *et* al (3) was used to identify significantly differentially expressed transcripts (FDR 5%) between the tumor and normal samples in each method, considering the two patients as replicates (since high dispersion was observed in the normalized signal values of Patient B adjacent normal sample with Method 2, in both the methods, patient A normal was used as reference for the tumor samples). Relevant reads, genes and regions of interest from Strand NGS were subsequently imported into GeneSpring 13.0 (Agilent Technologies) for integrated analysis with the proteome data. DE transcripts between the tumor and normal exhibiting a fold change ≥ 2.0 were used for all comparative analysis.

### Proteomics data: Protein identification and quantitation

Proteome Discoverer Beta Version 1.4 (Thermo Fisher Scientific Inc., Bremen, Germany) was used for the database searches. MS/MS data was searched using SEQUEST search algorithm against the RefSeq protein database (version 65). Trypsin was used as the enzyme with one missed cleavage allowed. Methionine oxidation was set as a variable modification, whereas methylthio of cysteine and iTRAQ modification at the N terminus of the peptide and lysine were set as static modifications. The tolerance for precursor and fragment mass was set to 20 ppm and 0.1 Da, respectively. The data was also searched against decoy database to calculate 1% FDR score as threshold at peptide level. Further, all the PSMs or peptides were selected using ‘high peptide confidence’ and ‘rank one’ peptide match filters in the Proteome Discoverer software and protein bias correction option was enabled for the normal distribution of the data. Protein identification with only unique peptide(s) was considered for further analysis. Relative quantitation of proteins was estimated based on the relative intensities of reporter ions released during MS/MS fragmentation of their peptides. Peptide data generated from the normal-tumor samples in each method were imported into Proteome Discoverer 1.4 software for protein and peptide identifications. All identified proteins were imported into GeneSpring 13.0. DE proteins (with ≥ 2 unique peptides) between the tumor and normal exhibiting a fold change ≥ 2 were used for all comparative analysis.

**Table S1:** **Coverage statistics (total number of aligned reads) for each sample.**

| Sample |  | Method 1 | Method 2 |
| --- | --- | --- | --- |
| Patient A | Normal | 42533366 | 75262608 |
|  | Tumor | 60193366 | 44041572 |
| Patient B | Normal | 32791046 | 25011674 |
|  | Tumor | 25238400 | 24901348 |

